# scRNA-seq revealed the rules for CDR3 length pairing in TCR beta and alpha chains and BCR heavy and light chains in human and mice

**DOI:** 10.1101/2024.05.28.596201

**Authors:** Jiaping Xiao, Jun Li, Yingjie Wu, Lanwei Zhu, Qi Peng, Yuanyuan Xu, Xinsheng Yao

## Abstract

The binding of T cell receptor (TCR)/B cell receptor (BCR) CDR3 with linear or conformational epitopes is a prerequisite for the adaptive immune response. The rules for CDR3 length pairing and the amino acid (AA) distributions in the prioritized rearrangement of the β chain (or heavy chain) and subsequent rearrangement of the α chain (or light chain) during the process of germline V(D)J gene rearrangement and self-tolerance selection to form the T- or B-cell CDR3 repertoire are currently unknown. Single-cell VDJ sequencing can provide abundant information on CDR3 sequences in paired chains. In this study, the paired-chain CDR3 sequences of central and peripheral T and B cells in humans and mice were analyzed according to single-cell VDJ sequencing data. This is the first study to find that T cells with β chains longer than paired α chains and B cells with heavy chains longer than paired light chains have absolute advantages. The proportion of T cells with length differences of three or more AAs between paired chains in CDR3 in humans was significantly higher than that in mice, and the proportion of B cells with length differences of six or more AAs between paired chains in CDR3 in humans was significantly higher than that in mice. The CDR3 length range of the β chain was narrower than that of the α chain, while the heavy chain had a much wider range than the light chain. The CDR3 length in human TCRs and BCRs was greater than that in mice. Extreme length differences were found between paired chains in both human and mouse T and B cells. There were significant differences in the range of pairing length, AA distribution, hydrophobicity, and polarity of the paired-chain CDR3 region in 5 chains (β, α, H, κ, and λ). Human TCR and BCR paired CDR3 sequences exhibited greater plasticity than those of mice. This study reveals new ideas about the molecular mechanisms of TCR and BCR responses to different antigenic epitopes. Our innovative findings offer novel perspectives and techniques for examining V(D)J rearrangement and selection mechanisms in T and B cells. These advancements enable us to analyze antigenic epitopes targeted by TCRs and BCRs, and further, to explore the genetic evolution of T and B cell responses across diverse animal species.

**Graphical Abstract:** scRNA-seq the TCR and BCR CDR3 repertoires of 35 samples from the central and peripheral tissues of humans and mice. It provides an opportunity to reveal the rules for CDR3 length pairing in TCR beta and alpha chains and BCR heavy and light chains.

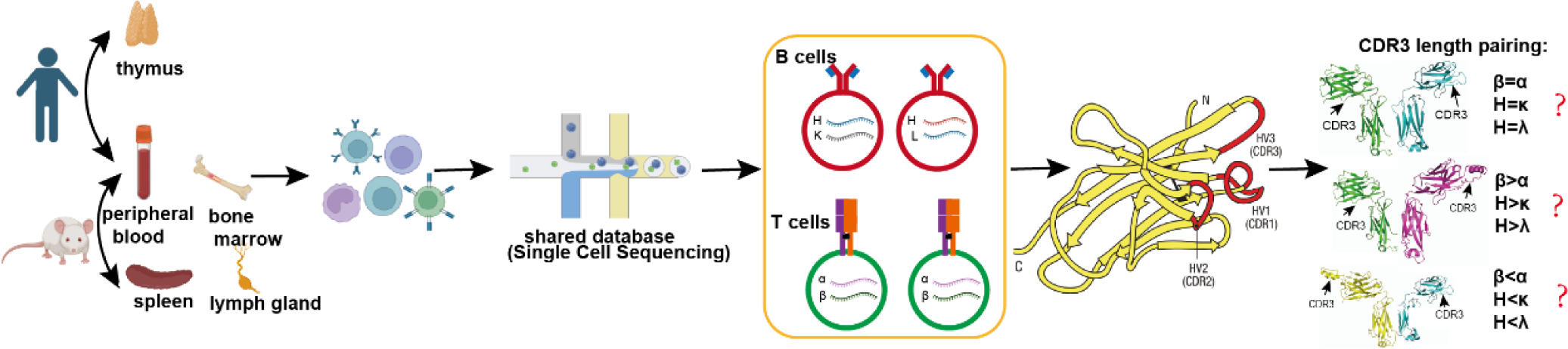

## Introduction

The αβ TCR is a dimer consisting of one α and one β chain, while the BCR is a tetramer consisting of two heavy chains and two light chains (κ and λ). The structural domain of the variable region in each TCR and BCR chain contains three complementarity-determining regions (CDRs) separated by four structurally conserved framework regions (FRs). CDR3 is a hypervariable region formed by V(D)J rearrangements and variable addition and subtraction of nucleotides at the junctions between gene segments^1^. CDR3 starts at 104-Cys in the germline V gene and includes positions 105 to 116 of the J gene (CDR3-IMGT has up to 12 amino acids), and positions 118-121 are conserved F/W-G-X-G motifs. In rearranged V-J or V-D-J genes and the corresponding cDNAs, CDR3 includes positions 105 to 117 (starting at the position preceding J-Phe or J-Trp), which exhibit gaps at the top of the loop in rearrangements of CDR3-IMGT less than 13 amino acids long or additional positions 112.1, 111.1, 112.2, 111.2, etc., in rearrangements of CDR3-IMGT greater than 13 amino acids long^2^.

The hypervariable region of CDR3 is the key site where T and B cells directly bind to the linear and conformational epitopes of antigens, and the length and amino acid distribution of CDR3 in paired polypeptide chains determine the breadth and depth of the adaptive immune response. The combination of paired α and β chains in CDR3 endows T cells with specificity for antigenic epitopes. The α chain and β chain CDR3 bind to the peptide-MHC molecule, and although the contribution of the β chain CDR3 binding to antigenic peptides is greater than that of the α chain CDR3, some structural analyses have shown that the α chain CDR3 also plays an important role^3^. BCR heavy chain and light chain CDR3 also participate in epitope binding simultaneously^4^. Studying the length, pairing rules, and AA distribution of CDR3 in two paired peptide chains is of great significance.

The range of CDR3 length varies greatly, as shown by the single-chain sequences obtained by high-throughput sequencing and spectral analysis of the CDR3 repertoire. Our previous studies indicated that there are 18 codon differences between the longest and shortest CDR3 sequences of the human TCR α chain and 19 codon differences in the TCR β chain^5^. In swine, there are 10 and 14 codon differences between the longest and shortest CDR3 sequences of the TCR α chain and β chain, respectively^6^. Moreover, the length of CDR3 in T or B cells among different species is also inconsistent. Ig in African Xenopus has 3-10 codon differences in the CDR3 sequence^7^, while humans typically have 5-12 codon differences. Our previous research also found that the BCR heavy chain CDR3 of humans is longer than that of mice^8^, and it is generally believed that mice have the shortest length, followed by humans, alpacas, Llama, and camels. The positive correlation between the length of the BCR heavy chain CDR3 and the body size of the species may be the basis for the differential response of B cells in different species^9^.

The length of CDR3 may be related to physiology or pathology. Differences in CDR3 length in the TCR β chain/BCR heavy chain were found among different age groups and different immune organs of the same species^10^. Under normal physiological conditions, CD4^+^ and CD8^+^ T cells of the thymus have relatively short CDR3 regions^11^, and the length of the BCR heavy chain CDR3 in elderly individuals is longer than that in younger individuals and longer in peripheral blood cells than in cells from the spleen^12^. However, Mangul S et al.^13^ found that tissue type has no effect on the length distribution of the CDR3 sequence of the BCR heavy and light chain in abundant human transcriptome data. Under pathological conditions, the immunoglobulin kappa light chains of rheumatoid arthritis (RA) expressed repertoires enriched for transcripts containing unusually long CDR3 lengths of 11 amino acid codons^14,15^, CDR3 length in CD45RA and CD45RO T-cell subpopulations related to infection is relatively long^16^, the TCR β chain CDR3 and BCR heavy chain CDR3 in patients with transplantation rejection have relatively shorter lengths^17^, IgA in nephropathy patients has a relatively short CDR3^18^, and the BCR heavy chain CDR3 length in effector B cells is longer in patients after SARS-COV-2 infection^19–21^.

The mechanism behind the differences in CDR3 length is related to the occurrence of different V/J gene families and rearrangement events (insertion and deletion)^4^. Fetal constraints on BCR CDR3 length begin with a genetic predilection for use of short DH/JH gene segments and through DH-specific limitations in N addition and deletion. Control of N addition may also exacerbate the block in B cell development^22^. In the same way, the Dβ sequence and N addition have been proven to control of TCR β chain CDR-B3 length, and the N addition and the sequence of the D is most likely to interact with the peptide that is bound to the presenting MHC^23^. Based on a structural simulation of TCR and antigen binding, Montemurro et al.^24^ proposed that accurate prediction of TCR and antigen binding models depends on paired α/β TCR sequences, and the specificity of the binding model for pMHC is relatively reduced only due to the single-chain CDR3 sequence. In addition, TCRs combined with the same epitope largely exhibit specific sequence and structural characteristics^25^. Sun X et al.^26^ analyzed 776 CDR3 sequences of human CD8^+^ T cells with specific responses and Kun Yu et al.^27^ found that the length range of CDR3 paired β and α chains was narrow, and there were differences in AA distribution.

The advent of single-cell high-throughput sequencing has revolutionized the analysis of T-cell receptor (TCR) and B-cell receptor (BCR) repertoires, enabling the examination of paired CDR3 sequence features from vast datasets. In this study, we harnessed this technology to comprehensively analyze the CDR3 repertoire across crucial anatomical compartments, including the thymus, bone marrow, spleen, and peripheral blood in humans and mice under physiological conditions. Moreover, we delved into the CDR3 repertoire of T and B cells targeting specific antigenic epitopes. Notably, our research represents the first in-depth analysis of TCR alpha and beta, as well as BCR heavy and light chain CDR3 length pairing features and rules. This unique approach aims to unravel novel techniques and research perspectives for understanding T and B cell V(D)J recombination mechanisms, the intricate interaction between TCR/BCR and antigenic epitopes, and the nuanced differences in T and B cell responses across diverse animal species. We anticipate that our findings will provide valuable insights into the complex dynamics of the immune system and facilitate the development of targeted immunotherapy strategies. (**Graphical Abstract**).

## Results

### T-cell CDR3 length analysis of paired TCR chains

A total of 26 single-cell sequencing samples were used for this analysis, including human central and peripheral αβ T cells and mouse peripheral and antigen-specific αβ T cells (**Table 1**). The CDR3 length distribution in all αβ T cells was found to follow a bell curve (**Supplementary Table 1**). In human central and peripheral T cells, the curve for the β chain exhibited peaks at 14 and 15 AA, and the α chain exhibited peaks at 13 and 14 AA (**Fig. 1A and B**). In mouse peripheral T cells, both α and β chains exhibited a peak of 14 AA (**Fig. 1C**). The distribution range of the length of the β chain CDR3 was narrower than that of the α chain in both humans (7-25 AA vs. 4-27 AA) and mice (6-25 AA vs. 4-25 AA). The average CDR3 lengths of the human α and β chains were 14.48±1.85 AA and 13.64±1.75 AA, both of which were significantly longer than those in mice (β chain: 13.73±1.44 AA; α chain: 13.59±1.35 AA, *P*<0.05).

**Fig.1.**
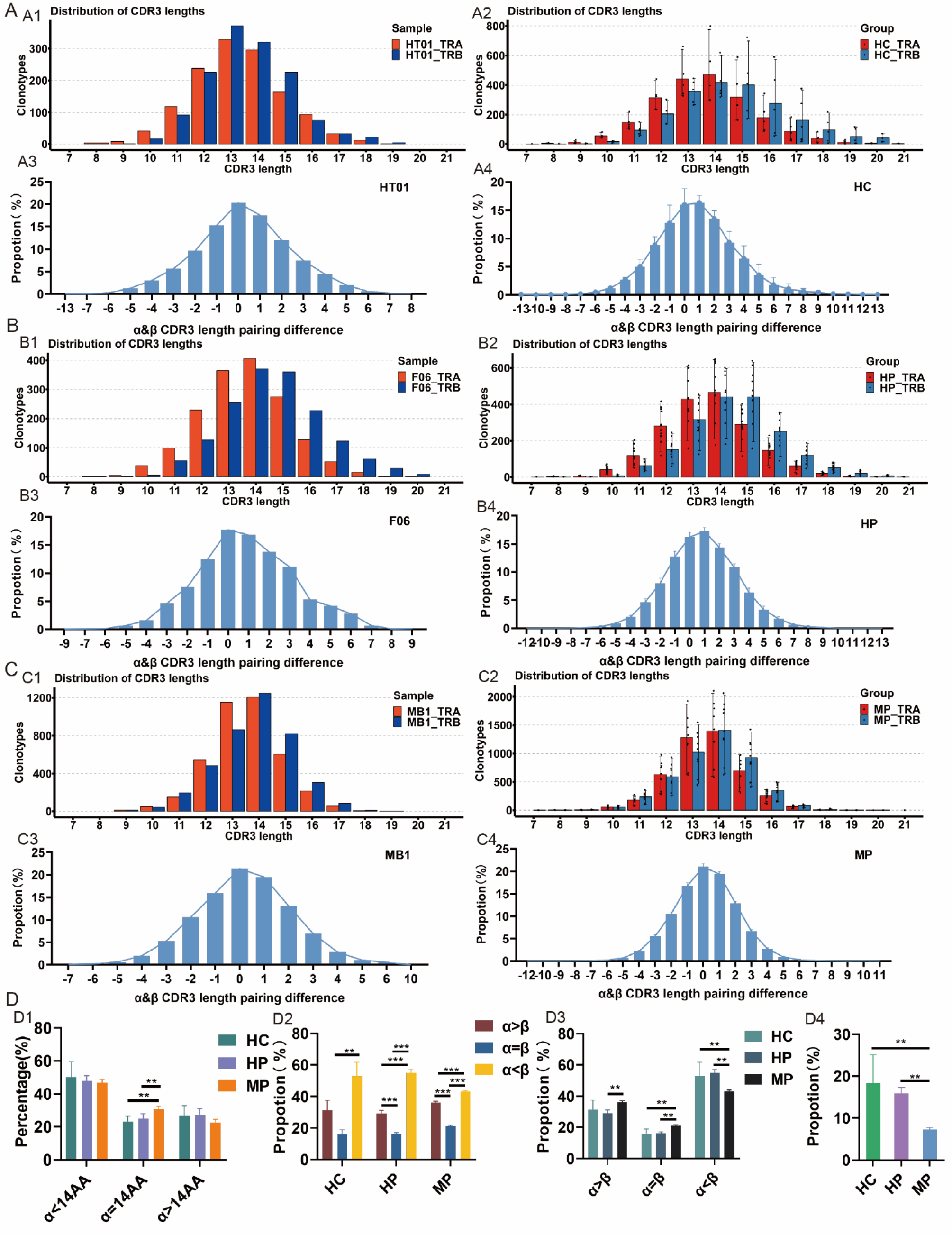
Analysis of CDR3 length distribution and pairing rule of TCR β and α chains. **(A) Human thymus samples (n=5)**. A1. CDR3 length distribution of β and α chains (HT01); A2. Statistics of the CDR3 length distribution of β and α chains in all five human thymus samples (HC); A3. CDR3 length pairing of β and α chains (HT01); A4. Statistics of the CDR3 length pairing of β and α chains in all five human thymus samples (HC). (**B**) **Human peripheral samples (n=12)**. B1. CDR3 length distribution of β and α chains (F06); B2. Statistics of the CDR3 length distribution of β and α chains in all 12 human peripheral samples; B3. CDR3 length pairing of β and α chains (F06); B4. Statistics of the CDR3 length pairing of β and α chains in all 12 human peripheral samples. (**C**) **Mouse peripheral samples (n=9)**. C1. CDR3 length distribution of β and α chains (MB1); C2. Statistics of the CDR3 length distribution of β and α chains in all nine mouse peripheral samples; C3. CDR3 length pairing of β and α chains (MB1); C4. Statistics of the CDR3 length pairing of β and α chains in all nine mouse peripheral samples. (**D**) **Intragroup and intergroup comparative analysis.** D1. The differences in α-chains paired with β-chains of 14AA in various TCR samples. E1. Intragroup comparative analysis; E2. Comparative analysis among groups; E3. Comparison of the proportion of paired chain CDR3 lengths with a difference of 3 AA (or more).

**Table.1.**
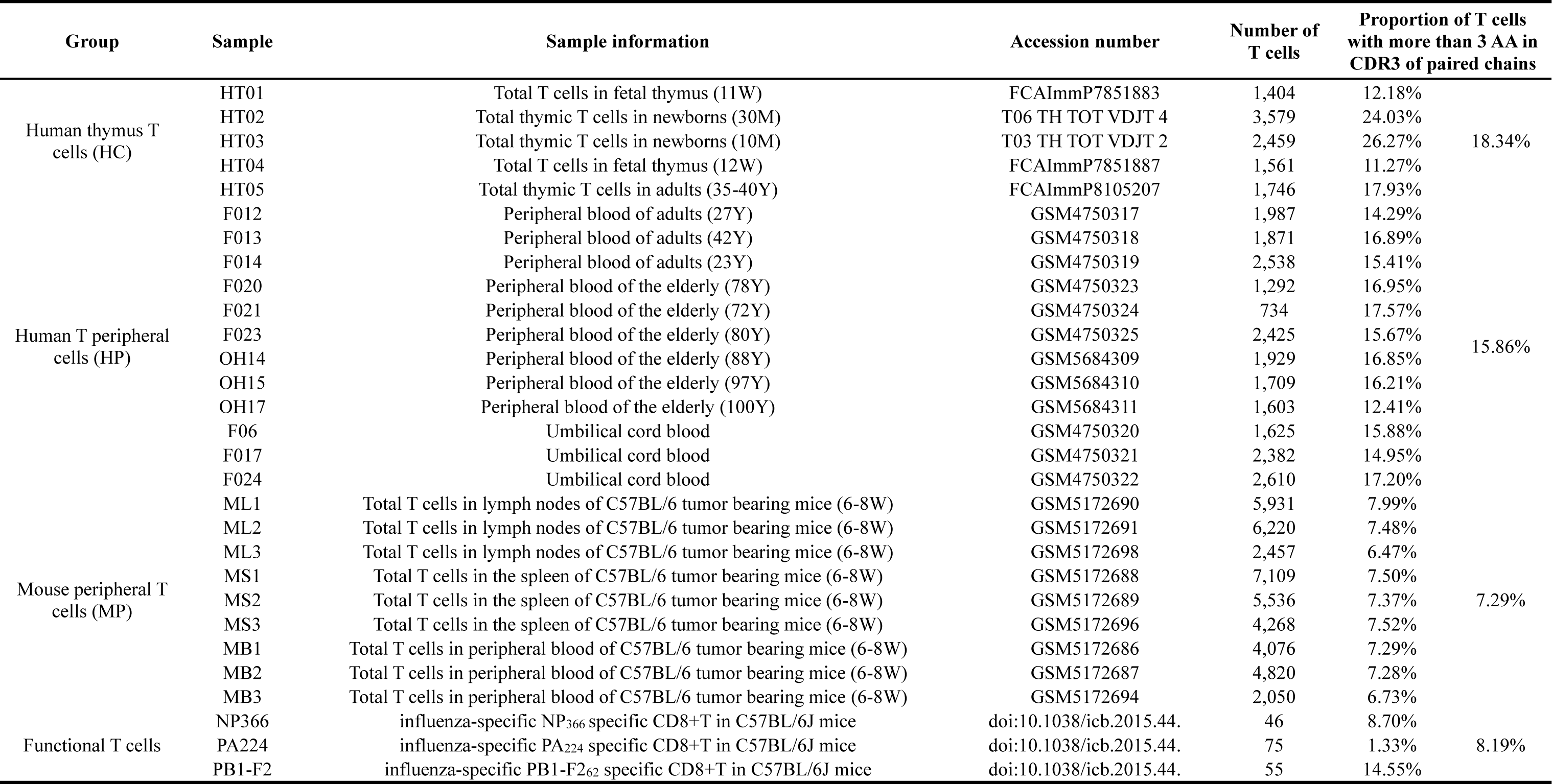
Single T cell sequencing data and statistical analysis of T cell with more than 3 AA in CDR3 region of paired chains.

By analyzing the differences in CDR3 length of two paired TCR chains (AA length of the β chain minus that of the paired α chain), it was found that the highest number of human central and peripheral T cells had one more AA in the β chain CDR3 region than in the paired α chain, and the differences exhibited a bell curve distribution ranging from −12 to 13 AA (**Fig. 1A and B**). Interestingly, the number of T cells with the same AA length in the two paired CDR3 chains was the highest in mice, showing a bell curve distribution with differences ranging from −12 to 11 AA (**Fig. 1C**). We selected T cells with a β-chain CDR3 length of 14 AA in the three groups and analyzed their paired α-chain CDR3 length. The α chain was found to have a bell-shaped distribution with peaks at 13 and 14 AA (range 4-22 AA) (**Fig. S1**). And the CDR3 length of mouse peripheral T cells, when paired with α- and β-chains both of 14AA, was significantly greater compared to those from human thymus (*P*=0.003) and human peripheral samples (*P*=0.002) (**Fig. 1D1**). Moreover, T cells with a longer β chain CDR3 compared to the paired α chain were dominant in all three groups, and the proportion of those T cells in human central and peripheral tissue was higher than that in mouse peripheral tissue (*P*<0.05). In contrast, the proportion of T cells with CDR3 lengths β=α and α>β in mice was higher than that in humans (*P*<0.01) (**Fig. 1D2 and D3**). The proportion of T cells with a difference of 3 AA or above between the paired β and α chains to the total number of αβ T cells was 18.34%, 15.86% and 7.29% in each of the three groups, respectively (**Table 1**), and the proportion of these T cells in humans was significantly higher than that in mice (**Fig. 1D4**).

### Antigen-specific CD8^+^T-cell, naïve and memory T-cell CDR3 length analysis of paired TCR chains

Three influenza-specific CD8^+^ T-cell lines were used for further analysis. Using the same method, we calculated the TCR CDR3 AA length difference between paired α and β chains. Effector CD8^+^ T cells from the influenza-specific NP_366_ and PA_224_ cell lines showed a longer (1-2 AA) α chain CDR3 length compared to the paired β chain. Effector CD8^+^ T cells from the influenza-specific PB1-F262 cell line showed the opposite result, represented as a bell curve distribution with β chain CDR3 longer than the α chain by one AA. The range of AA length differences of paired α and β chain CDR3 was −6 to 5 in the three influenza-specific CD8^+^ T-cell lines (**Fig. 2A**). The naïve and memory T cells were used for the further analysis. Just like the total T cells, in human naïve and memory T cells, the curve for the β chain exhibited peaks at 14 and 15 AA, and the α chain exhibited peaks at 13 and 14 AA. The distribution range of the length of the naïve and memory T cells β chain (10-20 AA vs. 10-21 AA) CDR3 was narrower than that of the α chain (7-27 AA vs. 7-19 AA). By analyzing the differences in CDR3 length of two paired TCR chains, it was found that the highest number of naïve T cells had one more AA in the β chain CDR3 region than in the paired α chain and the number of memory T cells with the same AA length in the two paired CDR3 chains was the highest. In memory T cells, the difference exhibited a distribution ranging from −6 to 8 AA, which is narrower than that of naïve T cells (−12 to 10 AA) (**Fig. 2B**).

**Fig2.**
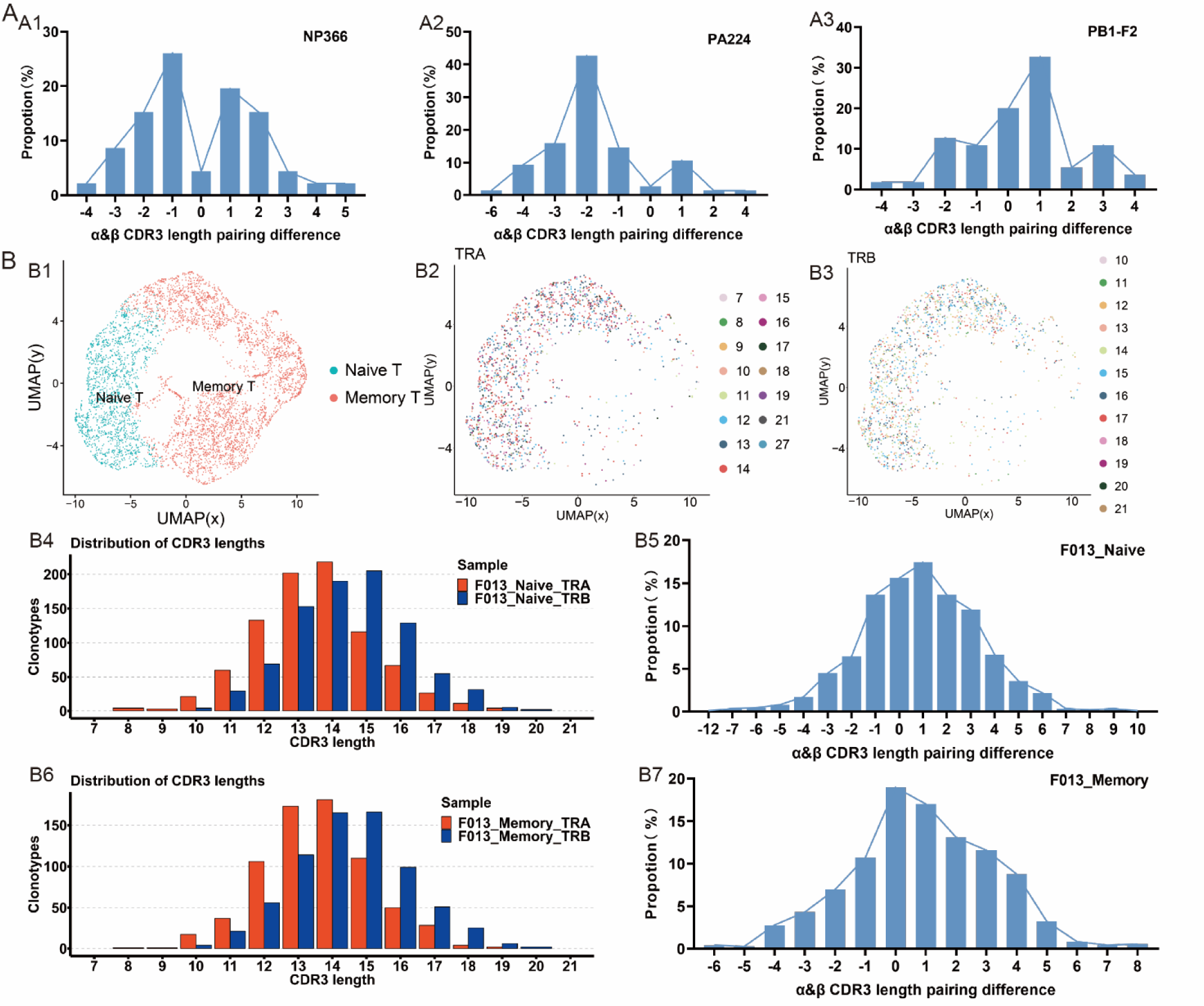
Analysis of CDR3 length distribution and pairing rule of Influenza-specific CD8+ T-cell, Naïve and memory T-cell TCR β and α chains. (A) Influenza-specific CD8+ T samples (n=3) A1. Statistics on the length difference of CDR3 paired chains (NP366); A2. Statistics on the length difference of CDR3 paired chains (PA224); A3. Statistics on the length difference of CDR3 paired chains (PB1-F2). (**B**) **Human peripheral blood naïve and memory T cells samples (F013).** B1. Clustering analysis of naïve and memory T cells; B2. Distribution of α chain CDR3 lengths used for clustering; B3. Distribution of β chain CDR3 lengths used for clustering; B4. CDR3 length distribution of β and α chains in naïve T cells; B5. CDR3 length pairing of β and α chains in naïve T cells; B6. CDR3 length distribution of β and α chains in naïve T cells in memory T cells; B7. CDR3 length pairing of β and α chains in memory T cells.

### T-cell CDR3 AA distribution of paired TCR chains

The AAs in human TCR β-chain CDR3 were mainly S, G, A, and F, and the α-chain mainly included G, A, F, and C. The AAs in mouse TCR β-chain CDR3 were mainly S, G, A, and F, and the α-chain mainly included A, G, S, and F (**Fig. 3A**). Human and mouse β chain CDR3 AA polarity scores were significantly higher than those of the paired α chain (**Fig. 3B**). The CDR3 AA polarity of human and mouse TCR paired α and β chains had no significant correlation (**Fig. 3C**). The hydrophobicity analysis of CDR3 AAs showed a significant correlation between the β chain and paired α chain in both human peripheral (*P*=0.012) and mouse peripheral tissues (*P*=0.024) (**Fig. 3D**).

**Fig.3.**
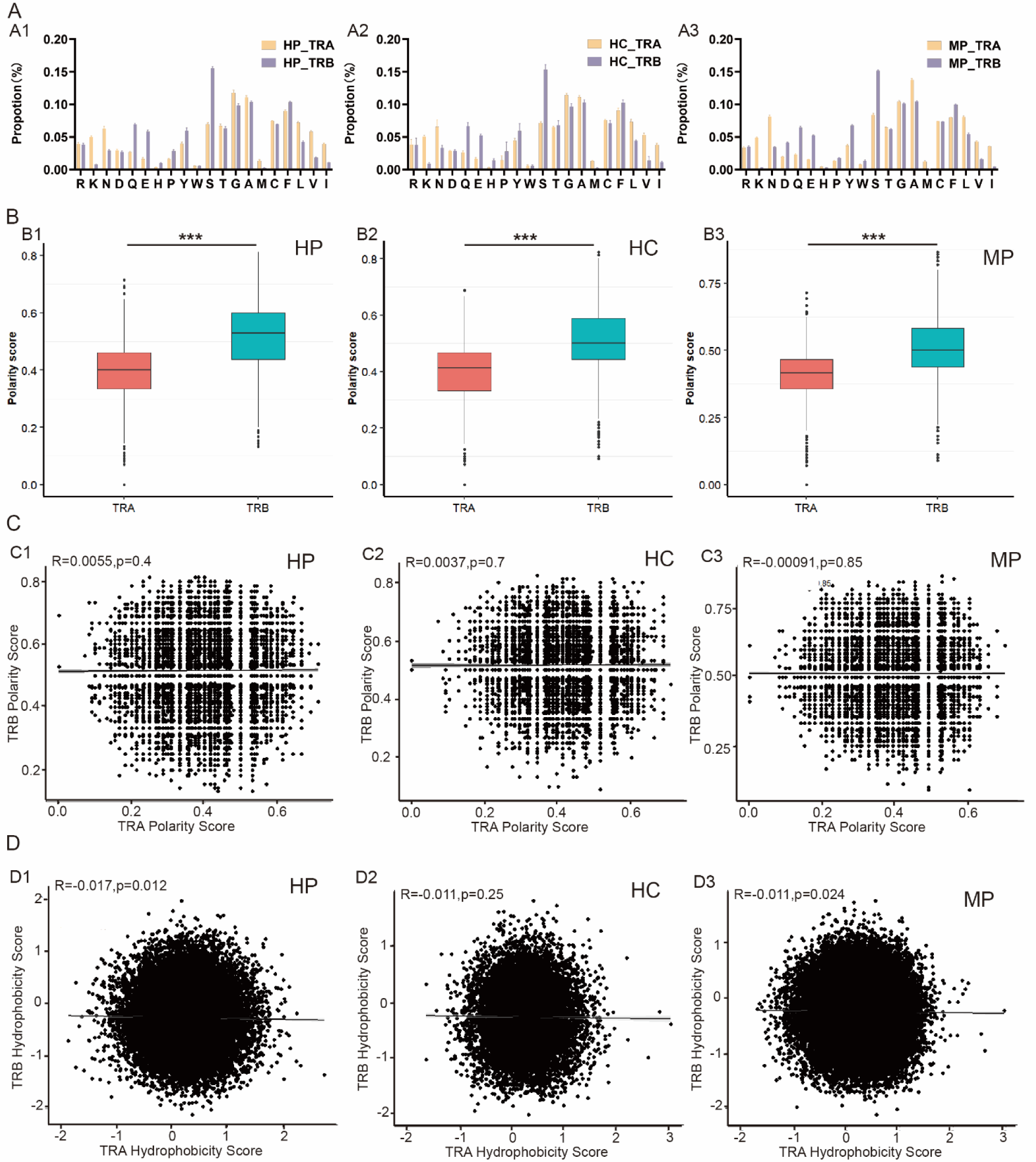
The characteristics of CDR3 AA in TCR paired chains. (A) The AA usage of CDR3 region. A1. Human peripheral samples (n=12); A2. Human thymus samples (n=5); A3. Mouse peripheral samples(n=9). (**B**) **The AA polarity of CDR3 region**. B1. Human peripheral samples(n=12); B2. Human thymus samples (n=5); B3. Mouse peripheral samples(n=9). (**C**) **Correlation of AA polarity in paired chains.** C1. Human peripheral samples(n=12); C2. Human thymus samples (n=5); C3. Mouse peripheral samples(n=9). (**D**) **Correlation of AA hydrophobicity in paired chains.** D1. Human peripheral samples(n=12); D2. Human thymus samples (n=5); D3. Mouse peripheral samples(n=9).

### B-cell CDR3 length analysis of paired BCR chains

A total of 9 single-cell sequencing samples were used for this analysis, including human and mouse central and peripheral B cells (**Table 2**). In human bone marrow and peripheral B cells (**Fig. 4A1, A3**), the curve of heavy chain length had a peak at 17 AA, which was 6 and 4 AA more than that of the κ and λ chains, respectively. In mouse bone marrow and peripheral B cells (**Fig. 4B1, B3**), the curves for both the κ and λ chains had peaks at 11 AA, which was shorter than that of the heavy chain (13 or 14 AA). The distribution range of both human and mouse heavy chain CDR3 length was wider than that of the κ and λ chains. The average CDR3 lengths of the human BCR heavy and λ chains were 4 and 2 AA longer than those of the mouse BCR heavy and λ chains, respectively. The CDR3 length distribution of the κ chain was consistent between humans and mice. The distributions of the CDR3 lengths of the κ and λ chains in humans and mice were biased (93.03%), which was significantly different from the β, α and heavy chains, which had a bell curve distribution (**Supplementary Table 1**). The lengths of human BCR CDR3 regions (heavy: 17.86±3.83; κ: 11.16±0.80 AA; λ: 12.57±1.08 AA) were generally significantly longer than that of mice (heavy: 13.72±2.30 AA; κ: 10.95±0.39 AA; λ: 11.35±1.13 AA).

**Fig.4.**
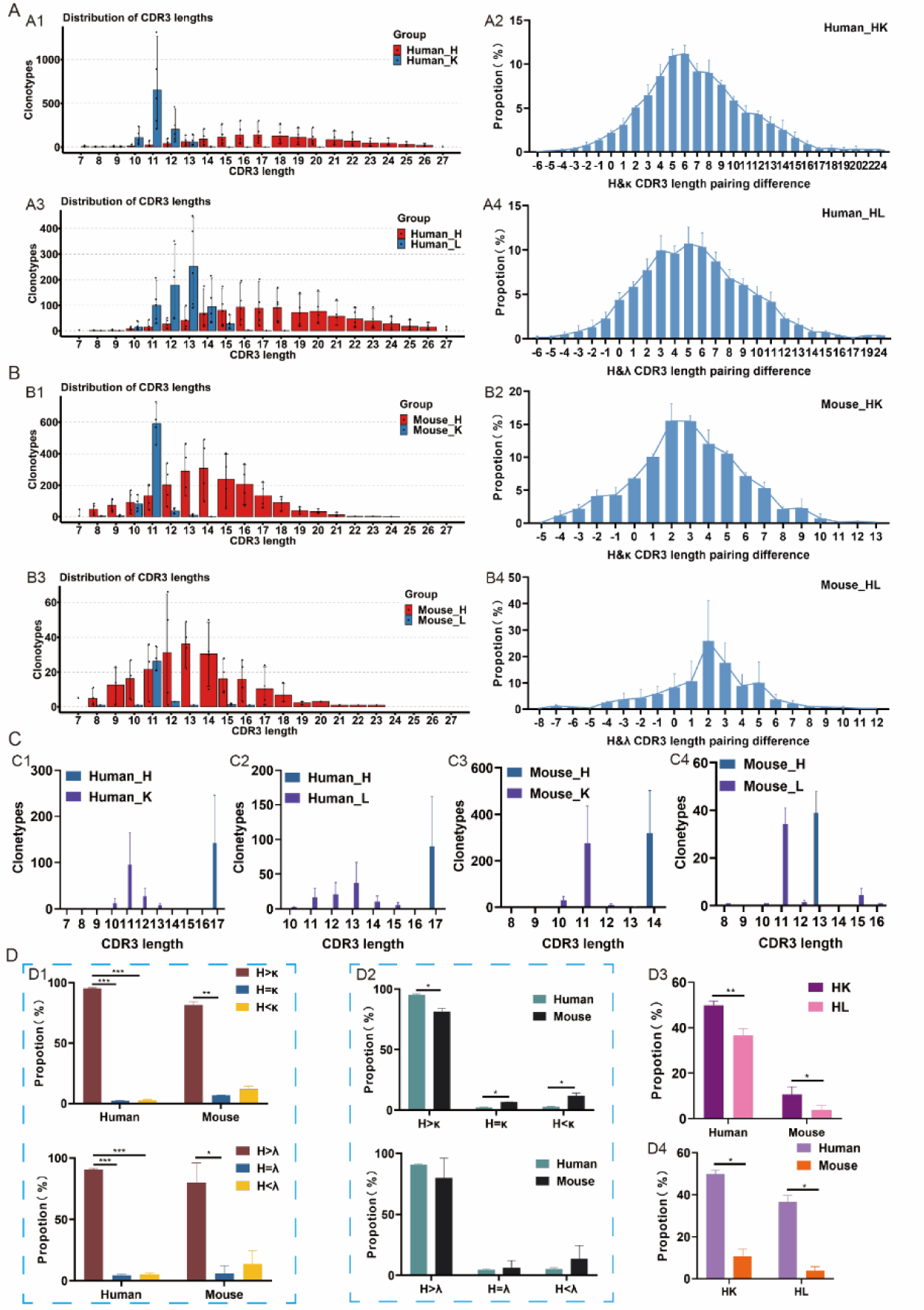
Analysis of CDR3 length distribution and pairing rule of BCR. **(A) Human bone marrow and peripheral samples (n=5)**. A1. Statistics of the CDR3 length distribution of H and κ chains; A2. the length difference of CDR3 in the BCR heavy chain and paired κ chain; A3. Statistics of the CDR3 length distribution of H and λ chains; A4. The length difference of CDR3 in the BCR heavy chain and paired λ chain. (**B**) **Mouse bone marrow and peripheral samples (n=4)**. B1. Statistics of the CDR3 length distribution of H and κ chains; B2. the length difference of CDR3 in the BCR heavy chain and paired κ chain; B3. Statistics of the CDR3 length distribution of H and λ chains; B4. The length difference of CDR3 in the BCR heavy chain and paired λ chain. (**C**) **Selection of paired κ or λ chain CDR3 length distribution rules when the highest peak of the BCR H chain CDR3 length is first rearranged**. C1. Human central and peripheral H chain CDR3 length = 17, paired κ-chain CDR3 length distribution (n=5); C2. Human central and peripheral H chain CDR3 length = 17, paired λ chain CDR3 length distribution (n=5); C3. Mouse central and peripheral H chain CDR3 length = 14, paired κ chain CDR3 length distribution (n=4); C4. Mouse central and peripheral H chain CDR3 length = 13, paired λ chain CDR3 length distribution (n=4). (**D**) **Intragroup and intergroup comparative analysis.** D1. Comparative analysis among groups; D2. Comparative analysis between humans and mice; D3. Comparison of the B-cell proportion of paired chain CDR3 lengths with a difference of 6 AA (or more) in human and mouse; D4. Comparison of the proportion of H and L (κ/λ) CDR3 length pairs differing by 6 AA (or more) between human and mouse.

By analyzing the differences in the AA length of BCR CDR3 paired chains (AA length of heavy chain minus that of the paired light chains), a bell curve distribution with a peak at 6/5 AA was found for the human central and peripheral BCR CDR3 length difference between heavy and paired κ/λ chain, ranging from −6 to 24 AA (**Fig. 4A2 and A4**). In mouse central and peripheral B cells, the difference in CDR3 length between heavy and paired κ/λ chains showed bell-like distributions, both with a peak at 2 AA, ranging from −5 to 13 AA and −8 to 12 AA, respectively (**Fig. 4B2 and B4**). We selected human H-chain CDR3 cells with a length of 17 AA and analyzed their paired κ- or λ-chain CDR3 lengths. We found that the κ-chain CDR3 length was 11 AA, and the λ-chain was distributed with 13 AA as the peak (range 10-15 AA). In the same way, we analyzed mouse B cells with H-chain CDR3 lengths of up to 13 or 14 AA and found that their paired κ and λ chains were both predominantly 11 AA (**Fig.4C**). The number of human B cells with heavy chain lengths longer than those of their paired light chains (κ and λ) dominated (Hκ, 95.22%; Hλ, 90.63%), significantly exceeding the number of B cells with heavy chain lengths ≤ paired light chain lengths. The pairing of heavy and light chains in four mouse samples showed the same pattern, with the highest number of B cells having heavy chains longer than those of paired light chains (Hκ, 81.42%; Hλ, 79.90%) (**Fig. 4D1**). The proportion of B cells with a longer heavy chain length than that of the κ chain was higher in humans than in mice (*P*<0.05), but the proportion of the other two types of B cells was higher in mice (**Fig. 4D2**). Focusing on B cells with CDR3 length differences greater than 6 AA between the paired chains (**Fig. 4D3**), we found that the proportion of B cells containing κ chains was significantly higher than that containing λ chains in both human and mouse samples (human, 49.75% vs. 36.53%; mouse, 10.64% vs. 3.88%), and the proportion of B cells with more than 6 AA between paired heavy and light (merged κ and λ) chain CDR3 sequences was higher in humans than in mice (*P*<0.05) (**Fig. 4D4**).

### Antigen-specific B-cell, naïve and memory B-cell CDR3 length analysis of paired BCR chains

SARS-CoV-2 RBD-specific B cells were used for further analysis. Using the same method, we calculated the difference in BCR CDR3 AA length between paired heavy and light chains (**Fig. 5A**). The CDR3 length distributions of heavy and light chains were 10-27 AA and 2-15 AA, respectively. More than 90% of antigen-specific B cells showed longer heavy chains than paired light chains, and the longest difference was 16 AA. A few antigen-specific B cells (<5%) showed longer light chains than paired heavy chains, and the differences were all 1 AA. The mouse central naïve and memory B cells were used for further analysis. In naïve B cells, the curve of heavy chain length had a peak at 14 or 13 AA, which was 3 and 2 AA more than that of the κ and λ chains, respectively. In memory B cells, the curves for both the κ and λ chains had peaks at 11 AA, which was 3 AA shorter than that of the heavy chain (15 or 14 AA). By analyzing the differences in the AA length of BCR CDR3 paired chains, the distribution with a peak at 3 AA was found for the naïve and memory B cells BCR CDR3 length difference between heavy and paired κ chains, ranging from −4 to 12 AA. Between heavy and paired λ chains, the distribution with a peak at 2 and 3 AA was found for the naïve and memory B cells, respectively. In memory B cells, the difference of heavy and paired λ chains exhibited a distribution ranging from −3 to 7 AA, which is narrower than that of naïve B cells (−7 to 10 AA) (**Fig. 5B**).

**Fig.5.**
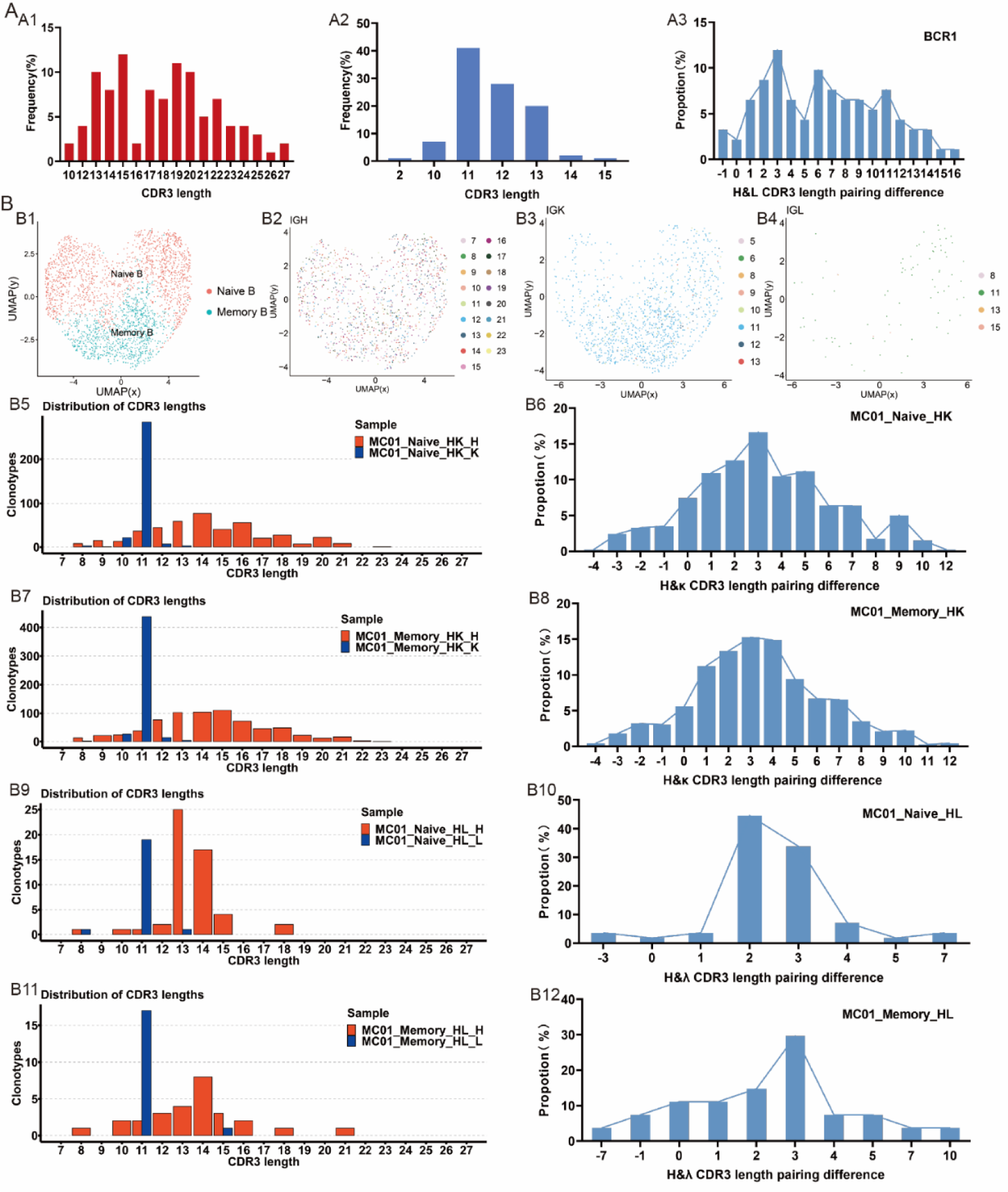
Analysis of CDR3 length distribution and pairing rule of RBD binding SARS-CoV-2 specific monoclonal antibody, Naïve and memory B-cell BCR. **(A) RBD binding SARS-CoV-2 specific monoclonal antibody**. A1. the length distribution of CDR3 in the BCR heavy chain; A2. the length distribution of CDR3 in the BCR light chain; A3. The length difference of CDR3 in BCR paired chains. **(B) Moue bone marrow naïve and memory B cell samples (MC01).** B1. Clustering analysis of naïve and memory B cells; B2. Distribution of H chain CDR3 lengths used for clustering; B3. Distribution of κ chain CDR3 lengths used for clustering; B4. Distribution of λ chain CDR3 lengths used for clustering; B5. CDR3 length distribution of H and κ chains in naïve B cells; B6. The length difference of CDR3 in the BCR heavy chain and paired κ chain in naïve B cells; B7. CDR3 length distribution of H and κ chains in memory B cells; B8. The length difference of CDR3 in the BCR heavy chain and paired κ chain in memory B cells; B9. CDR3 length distribution of H and λ chains in naïve B cells; B10. The length difference of CDR3 in the BCR heavy chain and paired λ chain in naïve B cells; B11. CDR3 length distribution of H and λ chains in memory B cells; B12. The length difference of CDR3 in the BCR heavy chain and paired λ chain in memory B cells.

### B-cell CDR3 AA distribution of paired BCR chains

We analyzed the AA distribution of BCR CDR3. In humans, the H chain AAs mainly include Y, G, A, and D, and the κ (or λ) chain AAs mainly include Q, T, F, and Y (or S, V, C, and F). In mice, the H chain AAs mainly include Y, A, W, and G, and the κ (or λ) chain AAs mainly include Q, T, F, and P (or W, Y, F, and C) (**Fig. 6A**). The heavy chain polarity scores of humans and mice were significantly lower than those of the paired κ chain, and the polarity score of the heavy chain was significantly higher than that of the paired λ chain in mice (**Fig. 6B**). There was no significant correlation of CDR3 polarity scores between heavy and paired light chains in either humans or mice (**Fig. 6C**). The CDR3 hydrophobicity score of the heavy chain was significant correlated (*P*<0.05) with the paired κ chain but not the paired λ chain (**Fig. 6D**).

**Fig.6.**
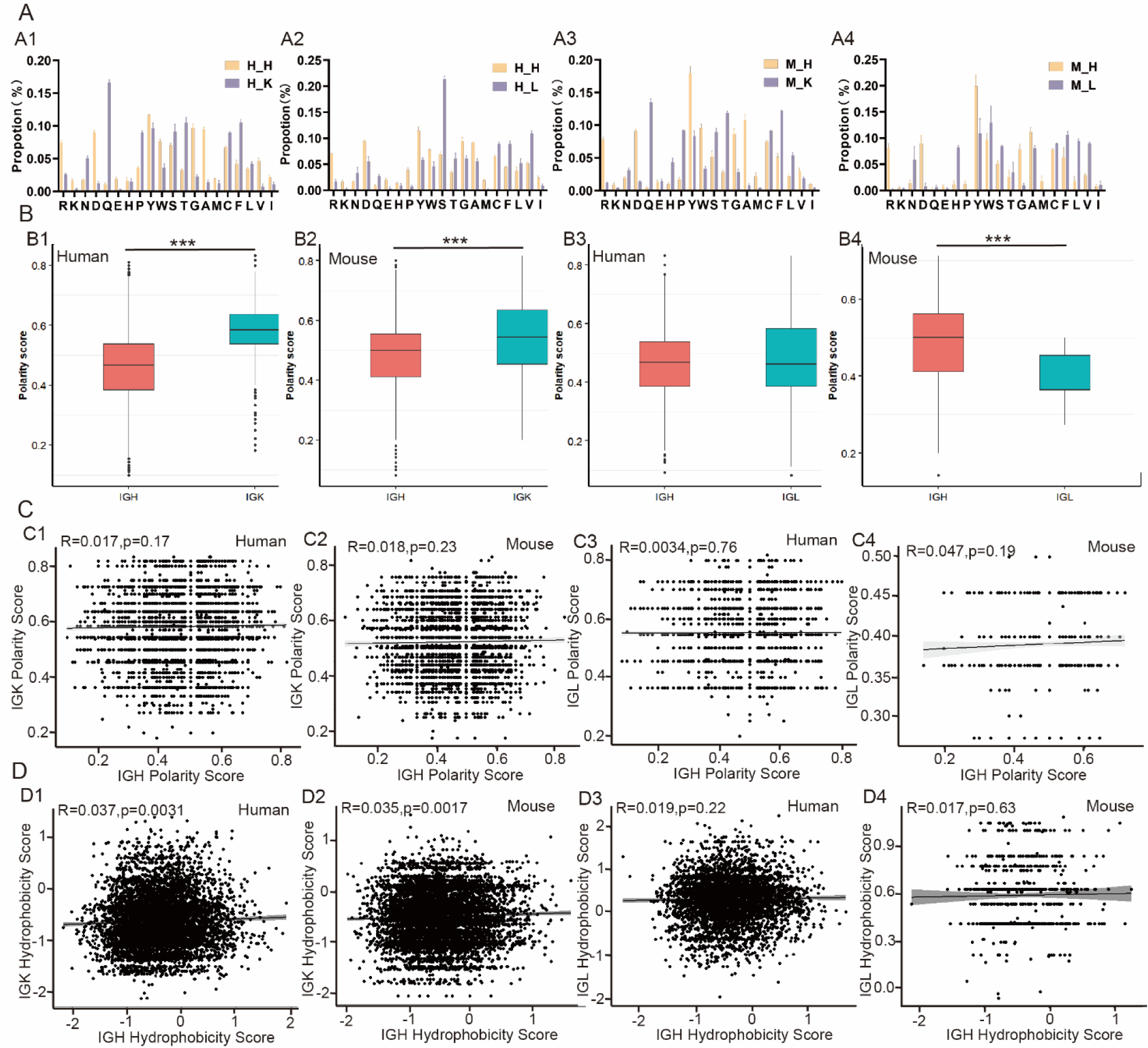
The characteristics of CDR3 AA in BCR paired chains. **(A) The AA distribution of the CDR3 region** A1. heavy chain and paired κ chain of humans (n=5); A2. heavy chain and paired λ chain of humans (n=5); A3 heavy chain and paired κ chain of mice (n=4); A4. heavy chain and paired λ chain of mice (n=4). (**B**) **The AA polarity of the CDR3 region.** B1. heavy chain and paired λ chain of humans (n=5); B2. heavy chain and paired κ chain of mice (n=4); B3. heavy chain and paired λ chain of humans (n=5); B4. heavy chain and paired λ chain of mice (n=4). (**C**) **Correlation of AA polarity in paired chains.** C1. heavy chain and paired λ chain of humans (n=5); C2. heavy chain and paired κ chain of mice (n=4); C3. heavy chain and paired λ chain of humans (n=5); C4. heavy chain and paired λ chain of mice (n=4). (**D**) **Correlation of AA hydrophobicity in paired chains.** D1. heavy chain and paired λ chain of humans (n=5); D2. heavy chain and paired κ chain of mice (n=4); D3. heavy chain and paired λ chain of humans (n=5); D4. heavy chain and paired λ chain of mice (n=4).

### Structure prediction of CDR3 with different AA lengths of paired chains

The T and B cells with the largest difference in CDR3 length between the paired chains were selected, and their CDR3 sequences were used for structural prediction. In the human TCR, the longest α chain was 26 AA, and the shortest paired β chain was 13 AA; the shortest α chain was 7 AA, and the longest paired β chain was 20 AA (**Fig. 7A; Fig.S2A**). In the mouse TCR, the longest α chain was 24 AA, and the shortest paired β chain was 12 AA; the shortest α chain was 12 AA, and the longest paired β chain was 23 AA (**Fig.7B**). In the influenza-specific paired TCR, the longest α chain was 14 AA, and the shortest paired β chain was 8-12 AA; the shortest α chain was 9 AA, and the longest paired β chain was 14 AA (**Fig. 7C**). In the human BCR, the longest κ (or λ) chain was 13 AA, and the shortest paired H chain was 7 AA; the shortest κ (or λ) chain was 7 AA, and the longest paired H chain was 27 AA (**Fig. 7D; Fig. S2A**). In the mouse BCR, the longest κ (or λ) chain was 11 AA, and the shortest paired H chain was 6 AA; the shortest κ (or λ) chain was 6 AA, and the longest paired H chain was 15 AA (**Fig.7E**). In the RBD-binding SARS-CoV-2-specific monoclonal antibody BCR sequence, the longest L chain was 11 AA, and the shortest paired H chain was 10 AA; the shortest L chain was 10 AA, and the longest paired H chain was 26 AA (**Fig. 7F**). The paired double chains of TCRs targeting specific antigenic epitopes only differ by 6 AA, and their CDR3 is relatively stable. There was a difference of 12 AA in the TCR paired chain CDR3 sequence of the initial human T cells, and CDR3 had stronger plasticity than that of mice. The structural prediction results of the CDR3 region in B cells were similar to those in T cells. The large length differences between paired chains in CDR3 provide greater flexibility for TCR and BCR to bind antigens.

**Fig.7.**
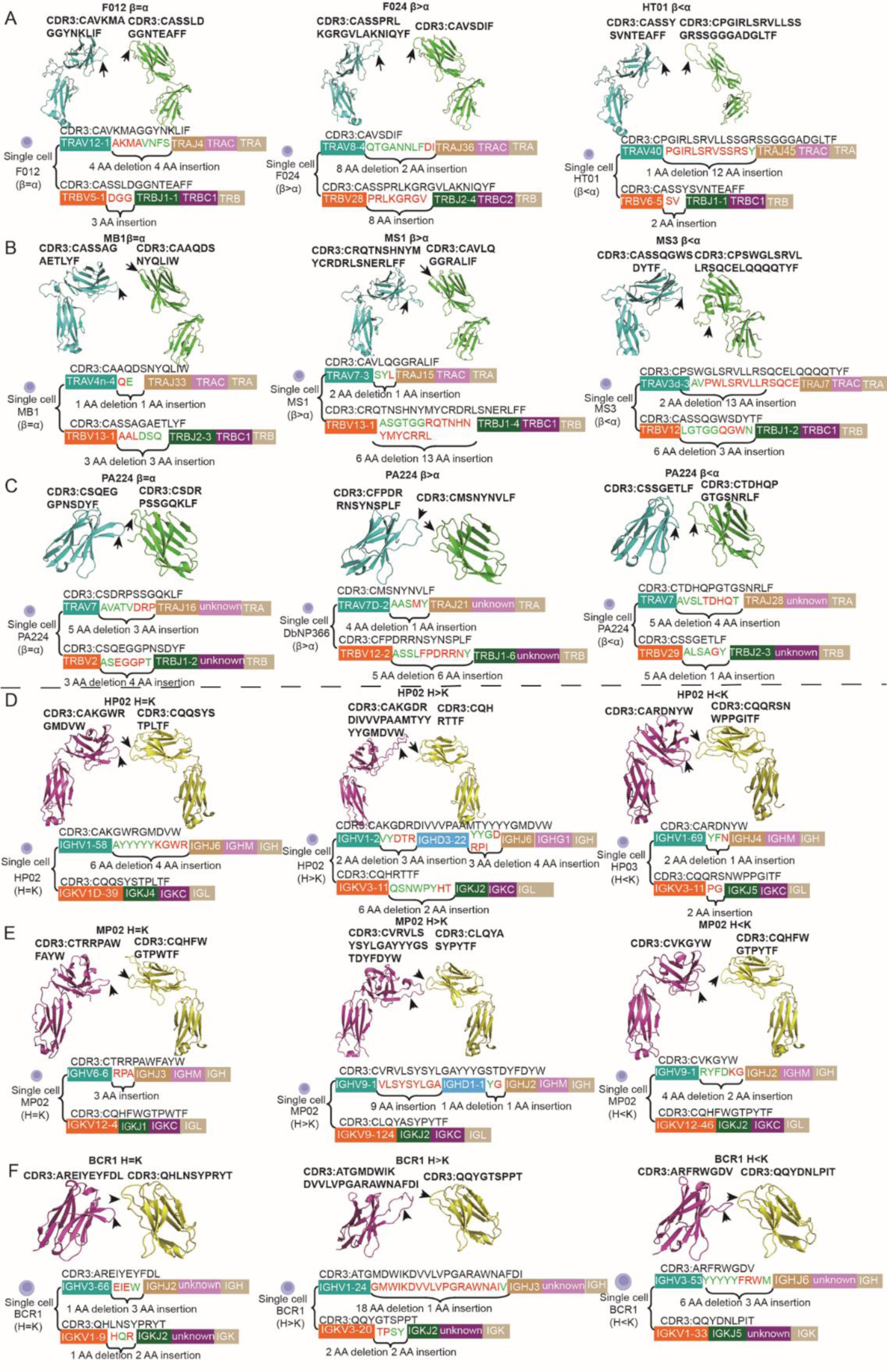
Examples of special paired sequences of TCR and BCR CDR3 and structure prediction. **(A)** The CDR3 sequence and structure prediction for human paired β and α chains with different lengths; (**B**) The CDR3 sequence and structure prediction for mouse paired β and α chains with different lengths; (**C**) The CDR3 sequence and structure prediction for paired β and α chains with different lengths in influenza-specific CD8+ T cells. (**D**) The CDR3 sequence and structure prediction for human paired heavy and κ chains with different lengths; (**E**) The CDR3 sequence and structure prediction for mouse paired heavy and κ chains with different lengths; (**F**) The CDR3 sequence and structure prediction for paired heavy and κ chains with different lengths in RBD binding SARS-CoV-2-specific monoclonal antibody.

## Discussion

Only a few paired chain sequences of TCR/BCR were obtained from individual T/B cells for multiplex PCR library construction and Sanger sequencing, which limited the study of the CDR3 length rules and characteristics of paired chains. With the rapid development of single-cell sequencing technology, single T/B-cell VDJ sequencing has yielded abundant paired CDR3 sequences. We utilized single T/B-cell VDJ sequencing data from the central and peripheral regions of humans and mice to provide a detailed explanation of CDR3 length rules and characteristics of two paired chains. The CDR3 length distributions of paired chains of human and mouse single T and B cells were bell curves with peaks at different lengths except for light chains, which is consistent with the length distribution results of single-chain CDR3 obtained through pedigree scanning, Sanger sequencing and HTS technology^5,8,28^. The CDR3 length of κ and λ chains exhibited irregular distributions with peaks at 13 AA and 11 AA, indicating that the V-J random rearrangement of the BCR light chain undergoes optimization during its pairing and tolerance selection process, resulting in a biased distribution. In the gene rearrangement model, the CDR3 lengths of the TCR beta chain and BCR heavy chain were close to a normal distribution, and it was proposed that the CDR3 length distribution may be related to optimization after rearrangement in addition to random selection during gene rearrangement^29^.

In analyzing the distribution range of CDR3 length, we found that the human TCR α chain CDR3 had a difference of 23 AA according to single-cell sequencing, which is longer than the difference of 18 AA found by our early CDR3 repertoire, while the human TCR β chain CDR3 had a difference of 18 AA, which is consistent with our early discovery of a difference of 19 AA^5^. The range of CDR3 length distribution in the human and mouse α chains was wider than that of β chains, which may be due to more than one allelic inclusion rearrangement of the VJ genes in the TCR α chain^30^, while the VDJ gene rearrangement in the β chain exhibits allelic exclusion. The CDR3 length range of the heavy chain in human and mouse B cells is wider than that of the paired κ and λ chains, which is consistent with the length of BCR CDR3 screened by transcriptome^13^. More amino acid insertions and cleavage occur during the rearrangement process of BCR heavy chain VDJ genes compared to the light chain, which may result in a longer heavy chain CDR3 sequence.

Both the TCR β chain and BCR heavy chain are formed after undergoing VDJ gene rearrangement, but the BCR heavy chain CDR3 has a wider length range, which may be related to the presence of more D genes with different lengths involved in the rearrangement (human, 7; mouse 6) compared to the TCR beta chain (both humans and mice have 2 D genes), as well as increased opportunities for nucleotide insertion. Although the V(D)J gene composition and rearrangement mechanism of TCR and BCR in humans and mice are basically consistent, we found that the CDR3 length range of human TCR and BCR chains is wider than that of mice, especially the heavy chain, indicating significant differences in the mechanism of overlength rearrangement between human and mouse TCR beta chains and BCR heavy chains. A BCR heavy chain CDR3 with more than 24 AA has been detected^31^, possibly due to an unconventional V (DD) J recombination^32–34^. The RSS before and after the IGHD gene is 9-12-7 and 7-12-9, and theoretically, there is no special recombination pattern of D-D. However, Safonova and Pevzner confirmed the 12/34 rule of D-D tandem rearrangement in IGH^35^, providing support for the formation mechanism of overlength CDR3. The overlength of TCR β chain CDR3 may also be related to the fusion rearrangement of TRBD1-TRBD2, in addition to the particularly large number of bases inserted during the V-D and D-J rearrangements^34^. In this study, we found significant differences in overlength CDR3 between humans and mice, which may be a potential mechanism for differences in adaptive response among different mammals. Moreover, we found that the human CDR3 (TCR β and α and BCR H and L) is longer than that of mice, which is consistent with the early findings of our team^8^ and other researchers^9^. The positive correlation trend between CDR3 length and species size is a very meaningful research direction, which may be closely related to the differential mechanisms of adaptive immune responses among different species.

In the CDR3 repertoire of individual total T and B cells, we innovatively discovered that CDR3 follows the main pairing rules of β chain length > α chain length and heavy chain length > light chain length, indicating that TCR β chains and BCR heavy chains may play a chief role in binding to a wide range of antigens. Our results are consistent with the structures of TCR and BCR^3,4^. The difference in CDR3 length between BCR paired chains is greater than that between TCR paired chains. This difference may be the basis for BCR and TCR binding to different antigenic epitopes. Our study provides new directions for research on the molecular mechanism of T- and B-cell binding to antigenic epitopes.

The flexibility of the CDR3 ring helps explain the cross-reactivity of TCR binding^36^. We found that the proportion of human TCR paired chain CDR3 sequences with a difference of 3 AA or more is significantly higher than that of mice, and the proportion of human BCR paired chain CDR3 sequences with a difference of 6 AA or more is significantly higher than that of mice, indicating that human TCR and BCR CDR3 have more structural flexibility than mice. Moreover, this result provides a new approach for studying the immune response differences to the same antigen between humans and mice and provides a basis for studying the unique immune responses possessed by other mammals from an evolutionary perspective.

CDR3 AA distribution in TCR and BCR paired chains affects the breadth and depth of their binding to antigenic epitopes. The conformational flexibility and amino acid composition of the TCR CDR3 have been reported to play an important role in TCR recognition and binding of antigens^37^. Early studies have shown that the frequency of glycine detection in the TCR β chain is higher than that in the TCR α chain^38^. This study was the first to analyze the CDR3 amino acid distribution of the central and peripheral TCR and BCR paired chains in humans and mice and found that both β and paired α chain CDR3 include G, A, and F, while BCR paired heavy and light chains exhibit significant differences. The TCR β chain CDR3 AA polarity scores of humans and mice were significantly higher than those of the α chain, and the CDR3 AA hydrophobicity of the β chain was significantly negatively correlated with that of the α chain. However, BCR CDR3 showed exactly the opposite results compared to TCR CDR3 in both CDR3 AA polarity scores and hydrophobicity, which may be the fundamental mechanism by which T and B cells respond to antigen epitopes. Stewart et al.^39^ proposed that the increase in length and the position dependence of glycine in TCR α and β chains may confer T cells the ability to cross-react and identify mutations or different pathogens. The research results of Borg et al. indicate that a single AA substitution in the CDR3 region plays an important role in the conformational and/or functional changes of TCRs, and the distribution of amino acid residues regulates the stability and/or flexibility of TCR binding recognition^40^. For Ig repertoire, the reading frame (RF)-specific sequence motifs of DH gene segments were enriched for tyrosine and depleted of hydrophobic and charged amino acids^41^, and progressively reducing CDR-H3 tyrosine content increasingly impairs preBCR checkpoint passage^42^. Moreover, with an accompanying increase in glycine, a shift in hydrophobicity was observed in the CDR-H3 loop from near neutral in fraction C (−0.08 +/− 0.03) to mild hydrophilic in fraction F (−0.17 +/− 0.02)^43^. The CDR3 AA distribution heterogeneity of TCR and BCR paired chains will provide insights into the molecular mechanisms of T- and B-cell responses to different epitope antigens.

Based on the abundant sequencing data of single T cells and single B cells, we found that there is a certain proportion of T and B cells in both humans and mice with significant differences in paired chain CDR3 length. This phenomenon may be related to specific antigenic epitopes, but its mechanism is currently unknown. For example, overlength BCR CDR3 produced widely neutralizing antibodies when targeting HIV-1 conserved epitopes in deep and obscured regions of the HIV-1 envelope^34,44^. Structural prediction of paired-chain CDR3 with significant differences suggested that the structures with the longest and shortest CDR3 paired chains have greater variability and flexibility in spatial conformation, which may be the basis for individuals to maintain their response to specific antigen epitopes. In antigen-specific T/B cells^45^, the maximum difference in CDR3 length between paired chains is 6 AA in functional CD8+ T cells, and antibodies targeting SARS-CoV-2 RBD epitopes^46^ also showed positive epitope binding ability and neutralizing effects, with a 16 AA difference in the BCR CDR3 length. However, Cukalac et al.^45^ proposed that analyzing a subset of cells or a single TCR chain cannot accurately describe the properties of the antigen-specific T-cell repertoire. Systematically conducting research on the source, mechanism, and significance of T and B cells with substantial length differences in CDR3 paired chains under individual physiological conditions, especially analyzing the crystal structure of extreme paired TCR and BCR chains, will be beneficial for special vaccine design and antibody therapy.

The pairing characteristics of two chains in a single T and B cell are the basis for studying their specific response function. The CDR3 length of the TCR and BCR paired chains and the dominant distribution and polarity of AAs are closely related to the function of T and B cells. Inappropriate pairing can lead to dysfunction of T and B cells^47^. The questions of how the α chain CDR3 length is selected to pair with the already rearranged β chain CDR3 and how the light chain CDR3 length is selected to pair with the already rearranged heavy chain CDR3 has been troubling immunologists. This study is the first to elaborate the CDR3 length distribution range, pairing rules, and AA composition characteristics of paired TCR and BCR chains and proposes that the plasticity of pairing in the human CDR3 region is significantly higher than that in mice.

In our study, it’s crucial to acknowledge certain limitations. Firstly, the length, amino acid composition, and average hydrophobicity of the receptor’s CDR3 that could potentially change during developmental stages^41,48,49^. These fluctuations might pose challenges to our understanding of receptor structure and function, hinting that cells might recognize antigens differently at various developmental or physiological states. Secondly, we must consider the differences existing among various immune cell subsets. These discrepancies could result in distinct receptor CDR3 lengths, amino acid compositions, or antigen recognition methods across different cell types^43,50^. Hence, further in-depth exploration of these variations is needed to comprehensively grasp their impact on immune system functionality.

## Methods

### scRNA-seq datasets of T cells

The scRNA-seq datasets of T cells were downloaded from NCBI, Gene Expression Omnibus (GEO) data repository and Zenodo. Total T cells of human thymus were derived from thymic tissue from the embryonic stage (9-39 weeks of gestation), neonatal period (0-28 days), and adult period (18-40 years old)^51^. Human peripheral T cells were derived from the cord blood of newborns, as well as peripheral blood mononuclear cells (PBMCs) from adult individuals (30.7±10.0 years old) and elderly individuals (85.8±11.1 years old)^52^. Mouse peripheral T cells were derived from the lymph nodes, spleen, and peripheral blood of C57BL/6 tumor-bearing mice at the age of 6-8 weeks^53^. Functional CD8^+^ T cells were derived from female C57BL/6J (H-2b) mice treated with HKx31 (H3N2) influenza virus^45^. The basic information of the above samples is shown in Table 1.

### scRNA-seq datasets of B cells

The scRNA-seq datasets of B cells were downloaded from the 10X Company, GEO and CoV-AbDab repositories. Among them, the human bone marrow samples (HC01) and mouse peripheral blood samples (MP01, MP02) were all provided by 10X Company. Human peripheral blood samples were obtained from healthy volunteers aged 41, 36, 68 and 65 years old^54,55^. Mouse bone marrow samples were derived from female C57BL/6 mice at the age of 8–12 weeks (accession number: E-MTAB-10174). RBD-based enzyme-linked immunosorbent assay (ELISA) screening resulted in the identification of 92 SARS-CoV-2–specific mAbs. The basic information of the above samples is shown in Table 2.

**Table.2.** Single B cell sequencing data and statistical analysis of B cell with more than 6 AA in CDR3 region of paired chains.

| Group | Sample | Sample information | Accession number | Number of B cells (Hk) | Proportion of B cells with more than 6 AA in CDR3 of paired chains |  | Number of B cells (Hλ) | Proportion of B cells with more than 6 AA in CDR3 of paired chains |  |
| --- | --- | --- | --- | --- | --- | --- | --- | --- | --- |
| Human B cells | HC01 | bone marrow | 10×company | 361 | 52.91% | 49.75 | 261 | 41.76% | 36.53% |
|  | HP01 | Peripheral blood (41Y) | GSM5550181 | 557 | 49.55% |  | 328 | 33.84% |  |
|  | HP02 | Peripheral blood (36Y) | GSM5550182 | 1,658 | 48.37% |  | 1,096 | 35.77% |  |
|  | HP03 | Peripheral blood (68Y) | GSM5550183 | 2,890 | 47.96% |  | 1,919 | 36.32% |  |
|  | HP04 | Peripheral blood (65Y) | GSM5831597 | 957 | 49.95% |  | 781 | 34.96% |  |
| Mouse B cells | MC01 | bone marrow of C57BL/6 mice (8-12W) | GSM4154732 | 1,439 | 15.01% | 10.64 | 112 | 5.36% | 3.88% |
|  | MC02 | bone marrow of C57BL/6 mice (8-12W) | GSM4154733 | 703 | 11.52% |  | 65 | 1.54% |  |
|  | MP01 | Peripheral blood of Balb/c mice | 10X company | 3,124 | 8.00% |  | 268 | 2.99% |  |
|  | MP02 | Peripheral blood of C57BL/6 mice | 10X company | 2,647 | 8.01% |  | 356 | 5.62% |  |
| Functional B cells | BCR1 | RBD binding SARS-CoV-2 specific monoclonal antibody | Sanjeev Kumar et al., 2022<br>(https://www.science.org/doi/10.1126/sciadv.add2032) | / | / |  | 92 | 56.52% |  |

### Screening of T and B cells

Off-machine data of single-cell sequencing may contain nonfunctional T/B cells. Therefore, we performed quality control on the downloaded data, including four items: “is_cell”, “high_confidence”, “type_chain” and “productive”. Cells with high confidence, expressing TCR chains and defined as functional (F) or nonfunctional (N) were used for subsequent analysis. Cells were classified according to the number of TCR chains expressed by a single cell. Then, combination analysis was performed according to the functionality of TCR chains. In this study, single αβ T cells containing only one functional TCR β chain and one functional TCR α chain were selected for analysis. The inclusion process for T cells was the same as for B cells, and single B cells containing only one functional BCR heavy chain and one functional BCR light chain (κ or λ) were selected for analysis.

### Cluster analysis of single-cell transcriptome data

The downloaded single-cell transcriptomic data was analyzed using R language (version 4.2.3) and the Seurat package (version 4.3.0). Initially, the dataset underwent quality control to eliminate doublets, dead cells, and other artifacts. Subsequently, the JackStraw and PCEIbow plotting functions were employed for principal component analysis (PCA). Selected principal components were then subjected to Uniform Manifold Approximation and Projection (UMAP) algorithm for dimensionality reduction and clustering analysis. Cell type annotations for the dataset were determined using the SingleR package (version 1.6.1) to identify specific cell types within each cluster. The T and B cell length information was integrated into the Seurat object using the AddMetaData function. Visualization of T and B cell subgroups’ CDR3 length information was carried out using the ggplot function.

### Analysis of CDR3 amino acid length for TCR and BCR paired chains

The preprocessed single-cell TCR and BCR paired chain sequences were saved separately in CSV format. The immunach package in R language was employed to output the CDR3 length distribution maps for the paired chains. The CDR3 amino acid lengths for the TCR and BCR paired chains were calculated using the length formula “=LEN()”. For TCR, the number of β chain CDR3 amino acids was subtracted from that of α chain CDR3 amino acids to obtain the difference in length. For BCR, the number of heavy chain CDR3 amino acids was subtracted from that of light chain (κ or λ) CDR3 amino acids to obtain the difference in length for the paired chains. The differences in the paired receptor chains for single T and B cells were counted and graphed using GraphPad Prism8.

### Characteristics of amino acid distribution in TCR and BCR paired chains

The distribution of amino acids in the paired chains was extracted using the formula “=LEN()-LEN(SUBSTITUTE(,“AA”,“”))” to count the 20 amino acids present in each sequence. The frequency of the amino acids was then calculated by the formula for each sample and graphed using GraphPad Prism8. R custom scripts were used to calculate the hydropathy and hydrophilicity scores for each sequence and to analyze the differences and correlations between the paired chains.

### Structure prediction of TCR and BCR with special paired sequences

The T and B cells with the maximum difference in CDR3 length in paired chains were selected, and the full-length amino acid sequences were filled in according to the V(D)J gene names (IMGT). AlphaFold2 software was used for structural prediction analysis. We also use the “multimer” model in Alphafold to generate the structural models of paired TCRs or Fabs of three samples.

### Statistical analyses

Data are presented as the mean ± SD. IBM SPSS Statistics 26 software was used for statistical analysis. The data conforming to a normal distribution were subjected to one-way ANOVA, and the rest were subjected to the Mann‒Whitney U test. *P*<0.05 was considered to indicate a statistically significant difference. Photoshop, R, Adobe Illustrator, and GraphPad Prism 8 software were used for data visualization.

## Data Availability

The scRNA-seq datasets of human central sample (HT01-HT05) were sourced from NCBI (SRA: ERP119282^56^) and can be accessed online through the Zenodo database (https://zenodo.org/records/5500511^57^)^51^.

The scRNA-seq datasets of human peripheral sample (F012,F013,F014, F020,F021,F023,OH14,OH15,OH17) were downloaded from GEO (accession number: GSE157007^58^)^52^.

The scRNA-seq datasets of mouse peripheral sample (ML1-ML3,MS1-MS3,MB1-MB3) were downloaded from GEO (accession number: GSE168944^59^)^53^.

Functional CD8+ T cells were derived from published^45^.

F013 single-cell transcriptome data were download from GEO (accession number: GSE157007^58^)^52^.

The scRNA-seq datasets of human central sample and mouse peripheral sample (HC01, MP01,MP02) were downloaded from 10×Company^60,61^.

The scRNA-seq datasets of human peripheral sample (HP01-HP04) were downloaded from GEO (accession number: GSE183051^62^; GSE194245^63^)^54,55^.

The scRNA-seq datasets of mouse central sample (MC01,MC02) were downloaded from NCBI (SRA: ERP127379^64^)^65^.

RBD-based enzyme-linked immunosorbent assay (ELISA) screening resulted in the identification of 92 SARS-CoV-2–specific mAbs were downloaded from CoV-AbDab^46^.

MC01 single-cell transcriptome data were download from GEO (accession number: GSE139836^66^).

All date files used to construct the figures, tables and statistical results are available from Zenodo repository^67^.

## Code Availability

This study does not use custom code.

## Acknowledgements

We thank to NCBI, GEO, 10×Company and Zenodo for the availability of the data. We thank to Park Luo, Bhatt, Shi, Z, Peterson, J.N, Wang P and Riedel R for providing shared the single-cell sequencing data.

## Author Contributions

Conceptualization: YXS

Methodology: XJP

Visualization: XJP, LJ

Funding acquisition: YXS

Project administration: XJP, YXS, LJ, WYJ, ZLW, PQ

Supervision: YXS, LJ

Writing – original draft: YXS, XJP, LJ, WYJ, ZLW, PQ

Writing – review & editing: YXS, XJP, LJ

## Ethics declarations

This study does not involve ethical requests or approvals.

