## supplementary Material for "scRNA-seq revealed the rules for CDR3 length pairing in TCR beta and alpha chains and BCR heavy and light chains in human and mice"

**Supplementary Materials**

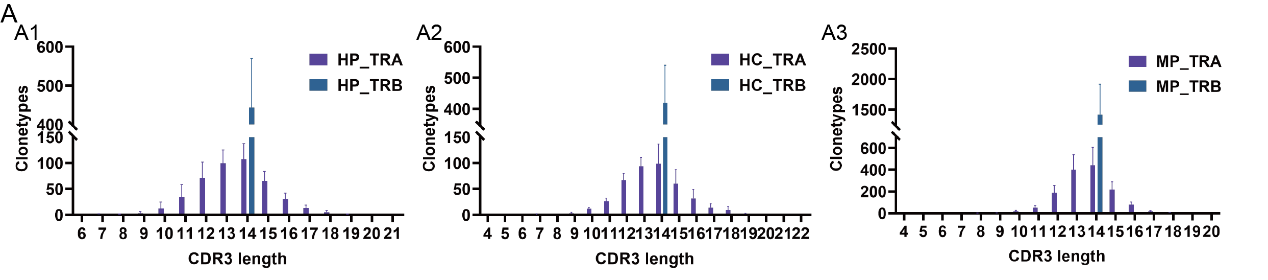

**Fig. S1.** **The rule that the highest CDR3 of the first rearranged β chain selects the paired α chain and the highest CDR3 of the first rearranged H chain selects the paired light chain**

A. **TCR first rearranged β chain CDR3 length peaks when selecting paired α chain length distribution rules:** A1. Human peripheral β chain CDR3 length=14, paired α-chain length distribution (n=12); A2. Human central β-chain CDR3 length=14, paired α-chain length distribution (n=5); A3. Mouse peripheral β chain CDR3 length=14, paired α-chain length distribution (n=9)

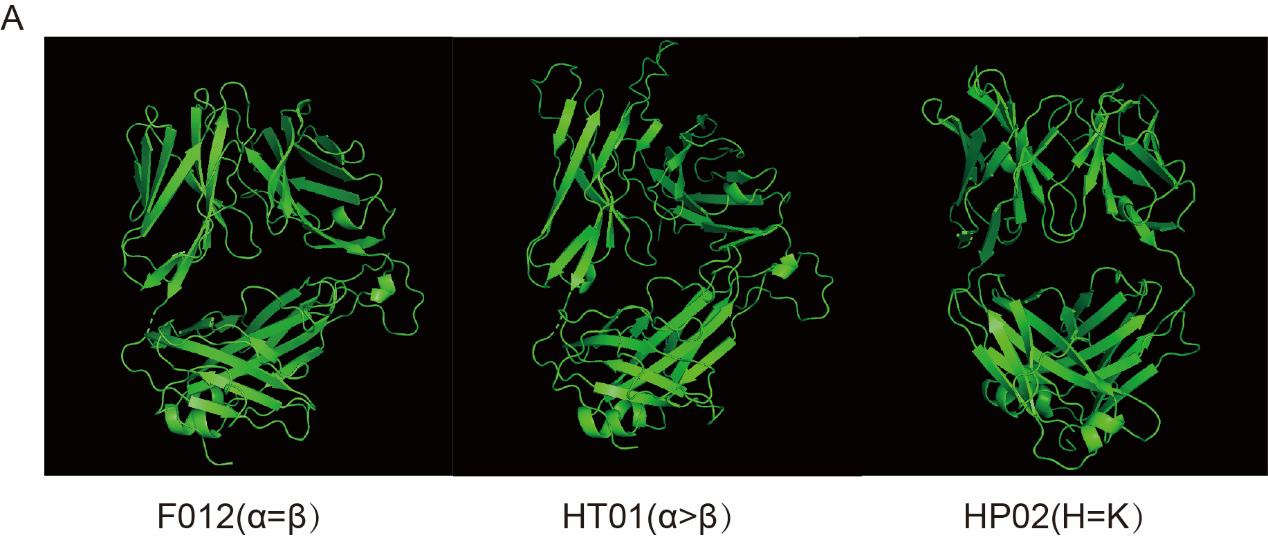

**Fig. S2.** **Use the "multimer" model in Alphafold to generate the structural models of paired TCRs or Fabs,** **The samples are labeled in order from left to right as F012(α=β), HT01(the longest α chain was 26 AA, and the shortest paired β chain was 13 AA), and HP02(H=K).**

**Supplementary Table 1：Calculation of normal distribution of each sample under 3 σ.**

| Sample | Chain | CDR3 length (μ-3σ, μ+3σ) | Actual distribution (%) |
| --- | --- | --- | --- |
| F012 | TRA | (8.53, 18.92) | 99.45% |
|  | TRB | (9.43, 19.22) | 99.60% |
| F013 | TRA | (8.35, 18.86) | 98.88% |
|  | TRB | (9.32, 19.65) | 99.57% |
| F014 | TRA | (8.53, 18.83) | 99.37% |
|  | TRB | (9.28, 19.66) | 99.37% |
| F020 | TRA | (8.68, 18.68) | 99.38% |
|  | TRB | (9.23, 19.79) | 99.54% |
| F021 | TRA | (8.45, 18.87) | 99.32% |
|  | TRB | (9.10, 19.70) | 99.05% |
| F023 | TRA | (8.66, 18.65) | 99.55% |
|  | TRB | (9.26, 19.57) | 99.34% |
| OH14 | TRA | (8.41, 19.05) | 99.17% |
|  | TRB | (9.46, 19.75) | 99.17% |
| OH15 | TRA | (8.74, 18.70) | 99.24% |
|  | TRB | (9.57, 19.59) | 99.47% |
| OH17 | TRA | (8.67, 18.64) | 99.44% |
|  | TRB | (9.58, 18.96) | 99.25% |
| F06 | TRA | (8.70, 18.63) | 99.63% |
|  | TRB | (9.16, 19.96) | 99.32% |
| F017 | TRA | (8.50, 18.38) | 99.45% |
|  | TRB | (9.16, 19.16) | 99.62% |
| F024 | TRA | (8.47, 18.43) | 99.46% |
|  | TRB | (9.19, 19.83) | 99.46% |
| T01 | TRA | (8.19, 18.46) | 99.64% |
|  | TRB | (8.88, 18.27) | 99.72% |
| T02 | TRA | (8.07, 19.72) | 99.25% |
|  | TRB | (8.65, 21.60) | 99.41% |
| T03 | TRA | (8.18, 19.47) | 99.35% |
|  | TRB | (8.64, 21.79) | 99.06% |
| T04 | TRA | (8.12, 18.13) | 99.55% |
|  | TRB | (8.74, 17.63) | 99.23% |
| T05 | TRA | (8.26, 19.30) | 99.43% |
|  | TRB | (9.01, 20.22) | 99.54% |
| MB1 | TRA | (9.57, 17.57) | 99.53% |
|  | TRB | (9.50, 18.05) | 99.71% |
| MB2 | TRA | (9.44, 17.73) | 99.29% |
|  | TRB | (9.36, 18.08) | 99.67% |
| MB3 | TRA | (9.68, 17.75) | 99.56% |
|  | TRB | (9.49,18.05) | 99.76% |
| ML1 | TRA | (9.54, 17.61) | 99.60% |
|  | TRB | (9.42, 18.07) | 99.60% |
| ML2 | TRA | (9.58, 17.64) | 99.68% |
|  | TRB | (9.41, 18.01) | 99.68% |
| ML3 | TRA | (9.48, 17.75) | 99.55% |
|  | TRB | (9.53, 17.93) | 99.51% |
| MS1 | TRA | (9.54, 17.56) | 99.56% |
|  | TRB | (9.36, 18.05) | 99.63% |
| MS2 | TRA | (9.59, 17.63) | 99.60% |
|  | TRB | (9.45, 18.01) | 99.69% |
| MS3 | TRA | (9.57, 17.65) | 99.67% |
|  | TRB | (9.41, 18.11) | 99.60% |
| HP01_HK | H | (7.03, 28.79) | 99.82% |
| HP01_HL | H | (7.48, 27.89) | 99.70% |
| HP02_HK | H | (6.46, 29.28) | 100.00% |
| HP02_HL | H | (6.61, 29.28) | 100.00% |
| HP03_HK | H | (6.02, 29.48) | 100.00% |
| HP03_HL | H | (6.35, 29.35) | 100.00% |
| HP04_HK | H | (6.11, 29.41) | 99.90% |
| HP04_HL | H | (7.19, 28.42) | 100.00% |
| HC01_HK | H | (5.65, 31.61) | 99.45% |
| HC01_HL | H | (6.21, 30.67) | 99.62% |
| MP01_HK | H | (5.41, 21.93) | 99.90% |
| MP01_HL | H | (4.81, 20.64) | 100.00% |
| MP02_HK | H | (5.21, 21.96) | 98.88% |
| MP02_HL | H | (4.84, 21.55) | 99.44% |
| MC01_HK | H | (5.54, 23.04) | 100.00% |
| MC01_HL | H | (7.91, 19.26) | 99.11% |
| MC02_HK | H | (5.52, 22.36) | 100.00% |
| MC02_HL | H | (9.17, 18.86) | 96.92% |
